# Effects of Biogas Slurry on Fruit Economic Traits and Soil Nutrients of *Camellia oleifera* Abel

**DOI:** 10.1101/472316

**Authors:** Lu You, Shuqin Yu, Huiyun Liu, Chutian Wang, Zengliang Zhou, Ling Zhang, Dongnan Hu

**Author notes:** Author’s Emails: Lu You, Shuqin Yu, HuiyunLiu, Chutian Wang, Zengliang Zhou, Ling Zhang, Dongnan Hu.

## Abstract

Soil nutrients play a principal role in *Camellia oleifera* Abel (oil-seed camellia) production. *Camellia oleifera* absorbs nutrients from surrounding soils and its production is highly influenced by nutrients or fertilization. In this study, we investigated the effects of biogas slurry applications on soil nutrients and economic traits of *C. oleifera* fruits. Five different amounts of fertilizing biogas slurry (0, 10, 20, 30, or 40 kg/plant/year from three applications per year) were applied to *C. oleifera* plants in 2015 and 2016. Rhizosphere soil nutrients and *C. oleifera* fruit economic traits (yield, seed rate, and oil yield)were measured. Fertilization with biogas slurryincreasedsoil organic matter, available nitrogen (N), phosphorus (P), and potassium (K) in both 2015 and 2016. Increases in soil available N, P, and Kwere largest at the highest slurry application rate and second largest at the second highest application rate. Fruit economic traits were maximized at the two highest application rates. Oil yield was correlated withsoil available P in 2015 and 2016, and soil organic matter in 2015. Fertilization with biogas slurry decreased saturated fatty acid content in fruit but had no effect on unsaturated fatty acid content. In conclusion, fertilization with biogas slurry increases rhizosphere soil nutrients and fruit economic traits of *C. oleifera* with the rates of at least 30 kg/plant/year having the most positive effects.

## Introduction

Biogas slurry is the secondary product of the anaerobic digestion process, and is also a widely used fertilizer in agricultural production. Biogas slurry is not only anenvironmentally friendly organic fertilizer, but it is also an efficient utilization of waste materials. Inrecent years in China, livestock wastes, such as feces and urine, havebecome a serious problem and create extensive environmental pollution [1], while anaerobic digestion is one of the effective solutions. The main product of anaerobic digestion is biogas which is an important and clean energy source. Meanwhile, the by-product, biogas slurry is also an environmentally friendly organic fertilizer thatcan be used in agricultural production [2–3]. Nowadays, the use of biogas slurry in agricultural production has drastically increased in China and many other Asian countries, not only because of the high cost of chemical fertilizers, but also for the high nutrients in biogas slurry [4–5]. It was reported that were more than 450 million tons of biogas slurry havebeen used in China each year [6]. Biogas slurry has dual effects in plant production. One is as a bio fertilizer with plentiful of nitrogen (N), phosphorus (P), potassium (K), and other trace elements, and the other is as a biological pesticide due tohigh levels ofamino acids, growth hormones, and antibiotics that promote plant growth [7–8]. Dry slurry was reported to contain <0.5% nitrogen, while the wet slurry contained 1.6% nitrogen as readily available nutrients. Because of the fermentation, ammonium ion (NH_4_^+^) content and pH of the biogas slurry increased, while the concentration of carbon (C) from the dry matter reduced, and the C/N ratio also decrease [9–10]. Furthermore, biogas slurry supplies more plant-readily available N than other fertilizers [11]. The available forms of nitrogen are inorganic, including nitrate (NO_3_) and ammonium (NH_4_) and simple structured organic partly from the degradation of organic matter. The available nitrogen can directly be absorbed by plants.

*Camelliaoleifera* C. Abel is an unusual oil shrub native to China that is distributed in 18 provinces/cities and more than 1,000 districts in China. It is reported that the planting area in China is over 5,000 acres, yielding about 200,000 tons of oil per year [12]. Camellia oil is a high quality edible oil, also named tea-oil, characterized byabundant unsaturated fatty acids, e.g., oleic acid and linoleic acid [13–14]. There is a long tradition of consumingtea-oil in China, especially in South China. In recent years, the planted area of *C. oleifera* is enlarging as demand for tea-oil is increasing [15]. The key factorthat determines the yield of *C. oleifera* is fertilizer application [16]. Traditional cultivation methodsdepend on chemical fertilizers, farm insecticides and chemical growth hormones which could lead to soil acidification, hardening, nutrientimbalances, and regression that results in reduction of production [17–19].

The effects of slurry and other organic manures on plants and crops have been shown[20]. Liquid fermented biogas slurry, from the outlet of the biogas digester can be readily used and applied directly to crops, vegetables, fodder grass and many other plants [21–23]. However, knowledge about the effects of biogas slurry on *C. oleifera* is limited. The potential benefits of biogas slurry to *C. oleifera* and its application at different amountsneed to be tested. In this study, we investigated the effects of biogas slurry applications on the soil nutrients and the fruit yield of *C. oleifera* to assess whether the usage of biogas slurry couldsubstitute partly or wholly for chemical fertilizers.

## Materials and methods

### Materials

This study was conducted in a *C. oleifera* plantation in Wannian, Jiangxi province in China from 2015 to 2016. The area has a typical warm and humid subtropical monsoon climate with an annual mean temperature of 17.4ºC, an annual rainfall of 1808 mm, and an annual relative humidity of 82%. The annual mean frost-free days are 259 d in the experimental area. The *C. oleifera* trees were planted in the red clay soil on the sunny hilly land with gradient less than 20%. The plantation in this study was seven years old, the row spacing was 3 by 3 m, and trees were 2-3 m high. The biogas slurry was prepared from pig farmyard manure using a farm biogas digester with a 200 m^3^ capacity. The biogas slurry was moderately alkaline, low in dry matter and ammonium.

### Experiment design

The experiment was carried out using a randomized block design with five treatments: (1) no biogas slurry [group B_0_]; (2) 10 kg of biogas slurry/plant/year [group B_1_] (3) 20 kg of biogas slurry/plant/year [group B_2_]; (4) 30 kg of biogas slurry/plant/year [group B_3_]; (5) 40 kg of biogas slurry/plant/year [group B_4_]. All of the five treatments were not fertilized by any chemical fertilizers. The biogas slurry was fertilized by the furrow method into the drip line of trees, divided into three times a year in March, June and September. Each treatment was carried out with three plots with 5 replicate plants in each plot.

### Detection methods

Mixed soil samples were collected from 0-20 cm and 20-40 cm rhizosphere soil immediately after fruit harvest. The concentrations of available N, P, K and organic matter were detected by using the methods of diffusion and absorption titration, NH_4_F-HCl extraction and molybdenum blue colorimetric, neutral NH_4_OAC extraction and flame photometric, and K_2_Cr_2_O_7_-H_2_SO_4_ and FeSO_4_ titration. Fruits of each experimental *C. oliefera* were harvested in October 2015 and 2016 and the weight of fresh fruit was measured. Because *C. oliefera* yield goes up and down each year [24–25], we calculated the production trait indices by using the average statistics of 2015 and 2016. Moisture rate of fresh seed= (fresh seed weight-dry seed weight)/fresh seed weight×100%, fresh seed rate = (fresh seed weight/fresh fruit weight) × 100%, dry seed rate = (dry seed weight/fresh fruit weight) × 100%, oil rate of kernel = (fat weight/kernel weight) ×100%, oil rate of fresh fruit = oil rate of kernel×dry seed rate×100%. Fatty acids in fresh fruit weremeasured according to the GB-5009,168-2016 method, by ShimadzuGC-2010 Plusgas chromatograph [26–27].

### Statistical analyses

The concentrations of organic matter, available N, P, and K between 2015 and 2016 were statisticallyanalyzed for significance by ANOVA by SPSS 19.0and means were compared by LSD (least significant difference) tests at p<0.05. The effects of biogas slurry on soil nutrients (organic matter, available N, P, and K) and yield were tested by correlation analysis.

## Results

### Effects of biogas slurry on organic matter in rhizosphere soil

Application of biogas slurry significantly increased the organic matter concentration of soils (Fig. 1A). In the first experimental year (2015), concentrations of soil organic matter raised as the dose of biogas slurry increased. Compared to the control group B_0_, the organic matter concentration of B_1_, B_2_, B_3_, and B_4_ groups increased by 32.2% (p<0.05), 55.8% (p<0.01), 70.9% (p<0.01), and 72.6% (p<0.01), respectively. Multiple comparison revealed no significant differences among treatments B_2_, B_3_ and B_4_. In the second experimental year (2016), the pattern was similar to that in 2015. All biogas slurry application rates increased organic matter compared the control group with the treatments B_3_ and B_4_ having higher enhancement rates than those in 2015, with increments of 142.28% and 137.56%, respectively.

**Fig.1.**
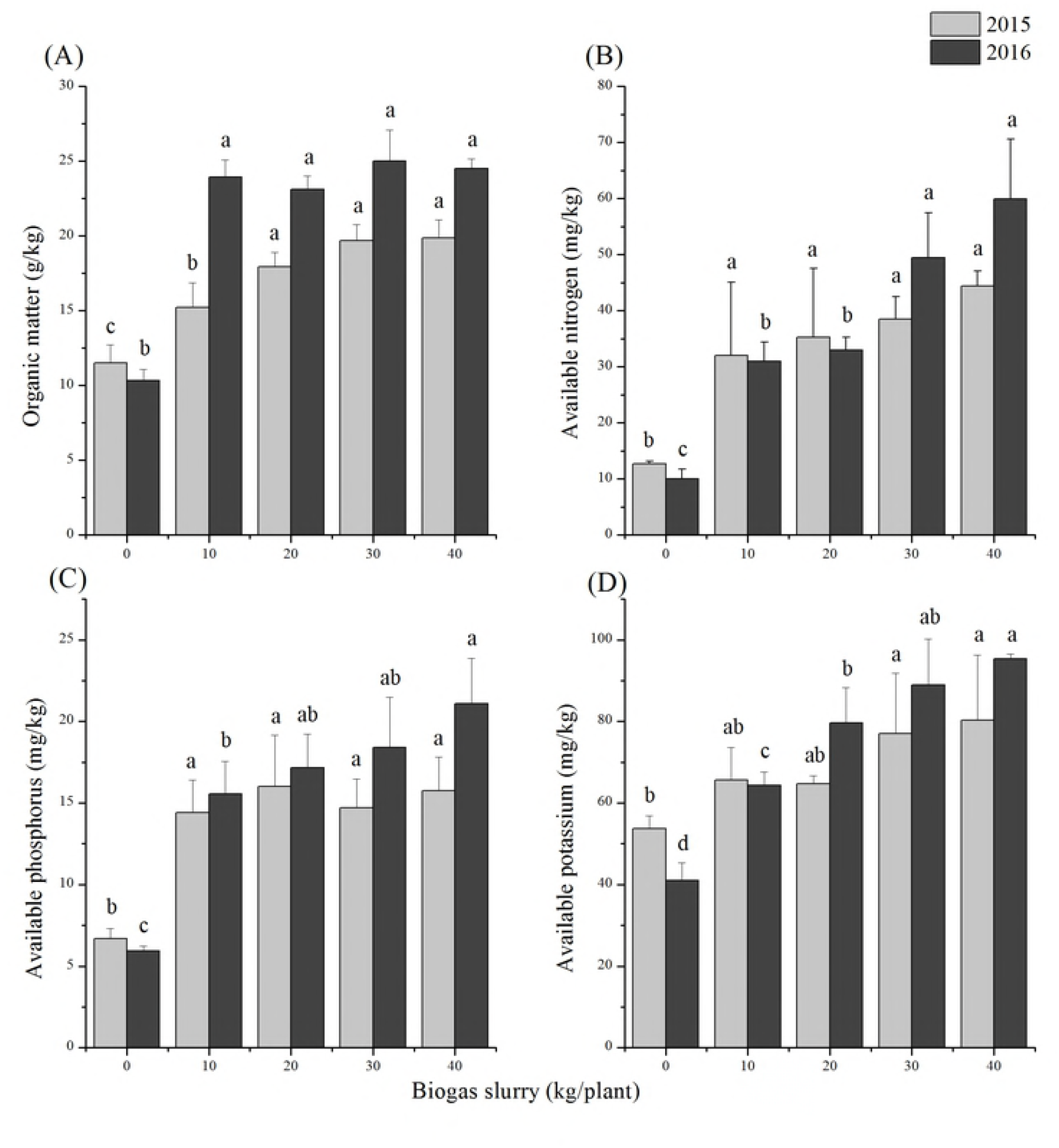
Effects of biogas slurry oncontents of (A) organic matter, (B) available nitrogen, (C) phosphorus and (D) potassium in rhizosphere soil during 2015 and 2016.

### Effects of biogas slurry on available nitrogen in rhizosphere soil

Fertilizing with biogas slurry increased soil available nitrogen both in the first (P_2015_=0.009<0.01) and second (P_2016_≈0.00<0.01) experimental years (Fig. 1B). Treatment B_4_, the highest biogas slurry dose used, caused the highest increament of available nitrogen in both 2015 (249.07%) and 2016 (499.2%), when compared to that in the control groups. Soil available nitrogen decreased from 2015 to 2016 in the no slurry addition control group. The lower application rates of slurry (B_1_, B_2_) had similar levels of soil available N in the two experimental years but at the second highest slurry addition rate (B_3_), soil available nitrogen continued to increase in the second year.

### Effects of biogas slurry on available phosphorus in rhizosphere soil

All four biogas slurry fertilized groups had higher concentrations of available phosphorus in 2016 than in 2015 (Fig. 1C). But, the control group had lower available phosphorus concentrations in 2016than in 2015. In 2015, the four fertilized groups (B_1_, B_2_, B_3_, B_4_) had increments of 151.81%, 139.97%, 119.96%, and 135.82% of available P, respectively, compared to the control group. Multiple comparisons revealed no significant difference among the four fertilized groups in 2015. In 2016, compared to the control group, the enhancement of available phosphorus was larger with greater slurry addition rates (B_1_, B_2_, B_3_, B_4_) with increments of 161.95%, 188.88%, 210.10%, and 255.05%, respectively.

### Effects of biogas slurry on available potassium in rhizosphere soil

Available potassium decreased from the first to the second year in the control and lowest slurry addition (B_1_) treatments (Fig. 1D). In 2015, available K in soils increased as the amount of biogas slurry application increased especially for treatments B_3_ and B_4_ (43.46% and 49.68%, respectively). In 2016, treatments B_3_ and B_4_ still showed significant enhancements of117.07% (p<0.01) and 132.52% (p<0.01), respectively.

### Effects of biogas slurry on fruit yield and main economic traits of *C. oleifera*

The average oil yield of *C. oleifera* in 2015 and 2016 showed a highest enhancement when fertilized with at least 30 kg biogas slurry per plant each year (Table 2). There was a trend of increasing yield as the amount of biogas slurry increased. Regarding to the main economic traits of *C. oleifera*, the fresh seeds from treatments B_0_ and B_1_ contained highest moisture ratios, while the lowest oil yield. Compared to the control treatment B_0_, the oil yield of treatments B_3_ and B_4_ increased in 105% and 95%, respectively.

**Table 1.** Experimental design (Unit: kg/plant)

| Treatments | March | June | September | Annual total |
| --- | --- | --- | --- | --- |
| B<sub>1</sub> | 0 | 0 | 0 | 0 |
| B<sub>2</sub> | 4 | 3 | 3 | 10 |
| B<sub>3</sub> | 8 | 6 | 6 | 20 |
| B<sub>4</sub> | 10 | 10 | 10 | 30 |
| B<sub>5</sub> | 14 | 13 | 13 | 40 |

**Table 2.**
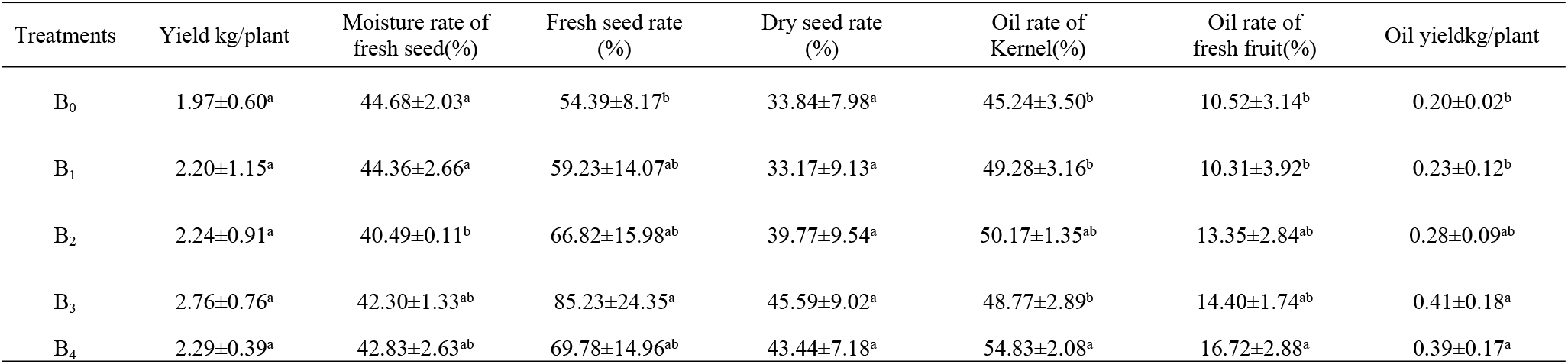
Effects of biogas slurry on yield and the main properties of *C. oleifera*

### Effects of biogas slurry on fatty acids of *C. oleifera* oil

Saturated fatty acids were mainly palmitic acid and stearic acid, accounting for about 10% of the fatty acid content (Table 3). The content of palmitic acid in the control B_0_ was the highest with all slurry application rates lowering the amounts. The effect of biogas slurry treatment on stearic acid was close to the significant level (P=0.07). Fertilizing with biogas slurry did not affect the unsaturated fatty acid content in fruit (Table 3). Correlations among saturated and unsaturated fatty acids showed that oleic acidwas negatively correlated with palmitic and linoleic acids, and linoleic acid was positively correlated with α-linolenic acid (Table 4).

**Table 3.** Saturated and unsaturated fatty acids in the fruit of *C. oleifera* fertilized by different amounts of biogas slurry

| Treatments | Saturated fatty acid % |  | Unsaturated fatty acid % |  |  |  |
| --- | --- | --- | --- | --- | --- | --- |
|  | Palmiticacid<br>(C16:0) | Stearic acid<br>(C18:0) | Oleic acid<br>(C18:1) | Linoleic acid<br>(C18:2) | γ- linolenic<br>acid(C18:3) | α-linolenic<br>acid(C18:3) |
| B<sub>0</sub> | 8.713±0.235<sup>a</sup> | 2.127±0.170<sup>a</sup> | 79.743±1.25<sup>a</sup> | 7.416±1.170<sup>a</sup> | 0.005±0.001<sup>a</sup> | 0.277±0.007<sup>ab</sup> |
| B<sub>1</sub> | 7.925±0.337<sup>b</sup> | 1.927±0.152<sup>ab</sup> | 78.695±0.574<sup>a</sup> | 7.804±0.485<sup>a</sup> | 0.007±0.001<sup>a</sup> | 0.252±0.017<sup>b</sup> |
| B<sub>2</sub> | 7.927±0.427<sup>b</sup> | 2.034±0.057<sup>ab</sup> | 79.620±0.984<sup>a</sup> | 7.286±0.284<sup>a</sup> | 0.005±0.002<sup>a</sup> | 0.260±0.013<sup>ab</sup> |
| B<sub>3</sub> | 8.009±0.436<sup>b</sup> | 1.820±0.059<sup>b</sup> | 79.674±1.629<sup>a</sup> | 7.695±1.308<sup>a</sup> | 0.004±0.002<sup>a</sup> | 0.301±0.033<sup>a</sup> |
| B<sub>4</sub> | 7.982±0.138<sup>b</sup> | 2.032±0.277<sup>ab</sup> | 80.616±0.485<sup>a</sup> | 6.533±0.511<sup>a</sup> | 0.005±0.000<sup>a</sup> | 0.241±0.031<sup>b</sup> |

**Table 4.**
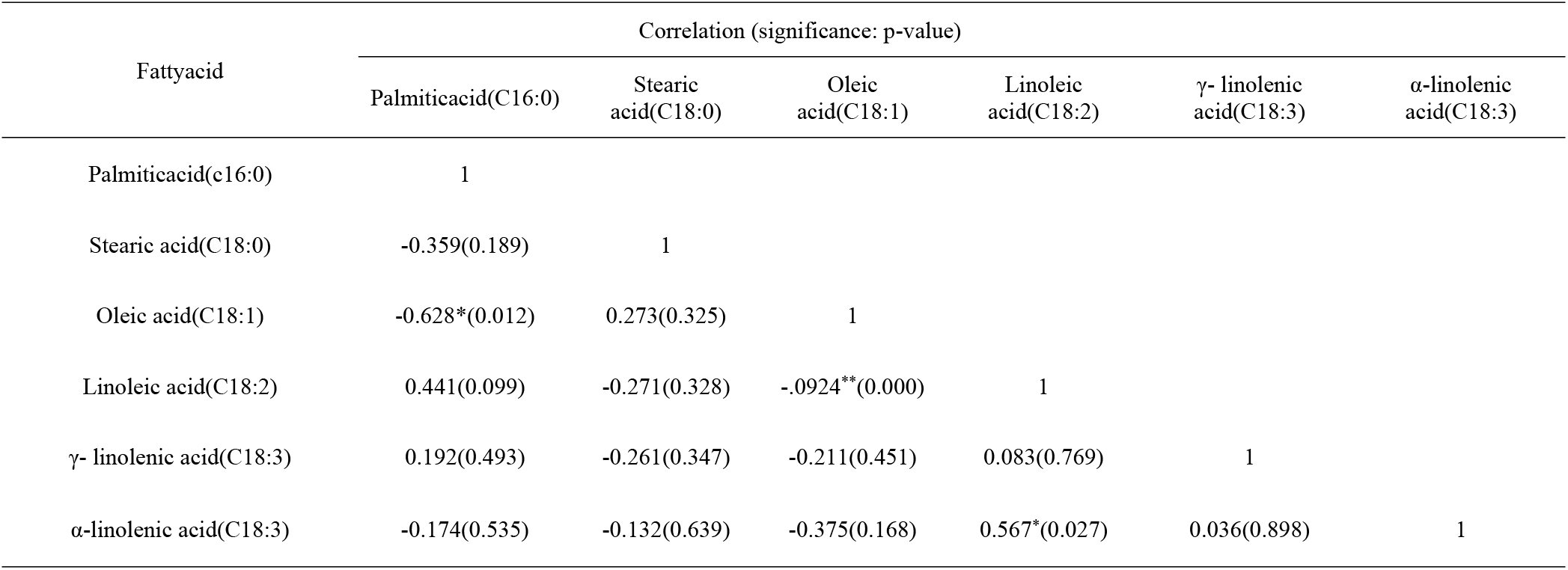
Correlations amongsaturated and unsaturated fatty acids

### Correlations among soil nutrients and fruit economic traits

The oil rates of fresh fruit were positively correlated with soil nutrients in 2015(N, P, K) and 2016 (organic matter; Table 5). Oil rates of kernels were positively correlated with N, P, K, and organic matter in both years (Table 5). Oil yield was positively correlated with the concentration of available P in both 2015 and 2016, and positively correlated with organic matter in 2015.

**Table 5.** Correlationsof oil yield components and soil nutrients

| Nutrients of soil | Year | Correlation (significance: p-value) |  |  |
| --- | --- | --- | --- | --- |
|  |  | Oil rate offresh fruit | Oil rate ofkernel | Oil yield |
| Available nitrogen | 2015 | 0.241(0.387) | 0.549*(0.034) | 0.407(0.132) |
|  | 2016 | 0.514*(0.050) | 0.720**(0.002) | 0.418(0.121) |
| Available phosphorus | 2015 | 0.472(0.076) | 0.628*(0.012) | 0.636*(0.011) |
|  | 2016 | 0.623*(0.013) | 0.681**(0.005) | 0.719**(0.003) |
| Available potassium | 2015 | 0.454(0.089) | 0.648**(0.009) | 0.252(0.364) |
|  | 2016 | 0.591*(0.020) | 0.647**(0.009) | 0.392(0.148) |
| Organic matter | 2015 | 0.681**(0.005) | 0.619*(0.014) | 0.644*(0.010) |
|  | 2016 | 0.372(0.172) | 0.565*(0.028) | 0.385(0.157) |

Note: \*indicates $p<0.05$ , $\geq 0.01$ ; \*\*indicates $p<0.01$ .

## Discussion

*Camellia oleifera* is a woody tree species endemic in China, and one of the important economic tree species. Oil from seeds is known to benefit health and is commonly eaten in China. Becauseflowers and fruits of *C. oleifera* grow throughout the year, and mature at the same time, fertilizers must be added to achieve high yields, especially of fruit and oil [28]. It is reported that P, N, K, Ca and Mg are the primary soil nutrientslimiting *C. oleifera* yield [29]. In conventional planting, chemical fertilizers are mainly used. But thelong-term useof chemical fertilizers hascaused land retirement, nutrient deficits and sealing of soil, which reduces yield [30]. To address this, new culture techniques and fertilizers should be developed and used in *C. oleifera* production. In this study, we used a new fertilizer, biogas slurry, in *C. oleifera* plantations, and investigated the response of nutrients in rhizosphere soil and the yields of *C. oleifera*.

Biogas slurry is a by-product of anaerobic fermentation, which plays a central role in the efforts to improve the utilization of animal manure and reduce the influence of animal excretion on surrounding environments [31–32]. During manuredigestion, about half of the carbon is released as methane and carbon dioxide (biogas), and part of the organic nitrogen is released as ammonium [33]. Ammonium is directly available to crops when it is appliedto fields. Furthermore, biogas slurry contains abundant available nitrogen, phosphorus and potassium, which are important nutrients for plants. It is reported that the supply of nitrogen from the digested slurry had a direct influence on the yield in the growing season, while the supply of phosphorus and potassium can be seen in the next year or several years [34]. So, we set a two-year experimental period to investigate the effects of biogas slurry on available N, P, and K of soils and the yield of *C. oleifera* in this study. During the two-year observation, we found the fertilization of biogas slurry has positive effects on the increment of available N, P, K of soils, and also improves the fruit and oil yield of *C. oleifera.* From the results in this study, we consider biogas slurry an effective substitute for chemical fertilizer in *C. oleifera* production.

Biogas slurry has abundant mineral elements and organic matter that can be released slowly. These characters of biogas slurry may positively affect soil fertility indices, e.g., organic matter, available N, P, and K over years [35]. A previous study evaluated the utilization ratio of NH_4_-N in biogas slurry, and found that more than 90% of the applied NH_4_-N could be used, which indicated an immediate increase in amount of the soil NH_4_-N [9]. Friedel *et al.* measured a 37% increase in inorganic N during the incubation of farmyard manure-derived biogas slurry in soil for 60 days [36]. Similarly, we observed a sharp enhancement of available N in soils fertilized with as little as 10 kg of biogas slurry per plant in 2015. The amounts of N supplied from some levels of biogas slurry application in this study were more than *C. oleifera* demand, so available N accumulation was observed in 2016 (Fig. 1B). Positive effects on available P and K after biogas slurry application were in accordance with that on available N, but with different increasing degrees. Available P is one of the main ecological factors limiting the increase of *C. oleifera* yield. Jun Yuan *et al.* studied responses to low P and found that *C. oleifera* roots secreted organic acids when the soil P is low and led to a utilization of soluble phosphates [37]. So, we just observed a slight but not sharp decrease of soil available P in the control group in 2016. Studies from Kashem *et al.* [38] demonstrated that alkaline pH could promote the availability of phosphorus in soils. It is likely that alkaline biogas slurry increased pH and, in turn, the availability of P in soils. The slow-release of nutrients in biogas slurry could contribute to the accumulation of organic matter, available N, P, and K in the second experimental year. We predict a larger promotion of nutrients in rhizosphere soil and yield of *C. oleifera* with long-term biogas slurry application.

## Conclusions

We studied the effects of biogas slurry on nutrients in rhizosphere soil and fruit economic traits of *C. oleifera.* We found that the use of biogas slurry could significantly enhance the concentration of available N, P, and K in soils, and significantly improve the yield of *C. oleifera.* In the first year, soils hadhigher concentrations of N, P, and K after treatment with biogas slurry and these enhancements were larger in the second year. Fertilized yield of *C. oleifera* oil also increased in the two experimental years, especially with higher application rates. Yield of oil also showed association to the increasing discipline of soil available N, P, and K in rhizosphere soils. Addition rates of at least 30 kg/plant/year (treatments B_3_ and B_4_) had the highest yield of fresh fruit, fresh seed rate, and dry seed rate, and resulted in a higher oil yield per plant. Because amounts beyond 30 kg biogas slurry per plant had no additional benefit to yield, so 30 kg biogas slurry per plantis likely the optimal rate of addition. We concludethatbiogas slurry has plays an important role in the production increasing of *C. oleifera*, and might be an effective substitution of chemical fertilizer in *C. oleifera* production.

## Acknowledgements

This study was financially supported by National Natural ScienceFoundation of China (grant number: 31560218, 41501317). The authors have no conflicts on this study and manuscript, and also appreciate Dr. Deping Song’s and Dr. Evan Siemann’s help inrevising the manuscript.

## Author Contributions

**Conceptualization:** Dongnan Hu

**Data curation:** Lu You, Huiyun Liu, Shuqin Yu

**Investigation:** Lu You, Chutian Wang, Zengliang Zhou

**Writing-original draft:** Lu You, Huiyun Liu, Ling Zhang

**Writing-review & editing:** Dongnan Hu, Lu You

## References

1. Zhang C, Su H, Baeyens J, Tan T (2014) Reviewing the anaerobic digestion of food waste for biogas production. Renewable and Sustainable Energy Reviews. 38: 383–392.

2. D’Imporzano G, Pilu R, Corno L, Adani F (2018) *Arundodonax* L. can substitute traditional energy crops for more efficient, environmentally-friendly production of biogas: A Life Cycle Assessment approach. Bioresource Technology 267: 249–256.

3. Holm-Nielsen JB, Al Seadi T, Oleskowicz-Popiel P (2009) The future of anaerobic digestion and biogas utilization. Bioresource Technology 100(22): 5478–5484.

4. Surendra KC, Takara D, Hashimoto AG, Khanal SK (2014) Biogas as a sustainable energy source for developing countries: Opportunities and challenges. Renewable and Sustainable Energy Reviews 31: 846–859.

5. Li JS, Duan N, Guo S, Shao L, Lin C, Wang JH, Hou J, Hou Y, Meng J, Han MY (2012) Renewable resource for agricultural ecosystem in China: Ecological benefit for biogas by-product for planting. Ecological Informatics 12: 101–110.

6. Mao C, Feng Y, Wang X, Ren G (2015) Review on research achievements of biogas from anaerobic digestion. Renewable and Sustainable Energy Reviews 45: 540–555.

7. Banik S, Nandi R (2004) Effect of supplementation of rice straw with biogas residual slurry manure on the yield, protein and mineral contents of oyster mushroom. Industrial Crops and Products 20(3): 311–319.

8. Insam H, Gómez-Brandón M, Ascher J (2015) Manure-based biogas fermentation residues – Friend or foe of soil fertility? Soil Biology and Biochemistry 84: 1–14.

9. Terhoeven-Urselmans T, Scheller E, Raubuch M, Ludwig B, Joergensen RG (2009) CO_2_evolution and N mineralization after biogas slurry application in the field and its yield effects on spring barley. Applied Soil Ecology 42(3): 297–302.

10. Hansen MN, Henriksen K, Sommer SG (2006) Observations of production and emission of greenhouse gases and ammonia during storage of solids separated from pig slurry: Effects of covering. Atmospheric Environment 40(22): 4172–4181.

11. Möller K, Müller T (2012) Effects of anaerobic digestion on digestate nutrient availability and crop growth: A review. Engineering in Life Sciences 12(3): 242–257.

12. Liang HY, Hao BQ, Chen GC, Ye H, Ma J (2017) Camellia as an Oilseed Crop. HortScience 52(4): 488–497.

13. Ma J L, Ye H, Rui Y K, Chen GC, Zhang N (2011) Fatty acid composition of *Camellia oleifera* oil. Journal fürVerbraucherschutz und Lebensmittelsicherheit 6(1): 9–12.

14. Zhu XY, Lin HM, Chen X, Xie J, Wang P (2011) Mechanochemical-Assisted Extraction and Antioxidant Activities of Kaempferol Glycosides from *Camellia oleifera* Abel. Meal. Journal of Agricultural and Food Chemistry 59(8): 3986–3993.

15. Wang XN, Chen YZ, Wu LQ, Liu RK, Yang XH, Wang R, Yang KW (2008) Oil Content and Fatty Acid Composition of *Camellia oleifera* Seed. Journal of Central South University of Forestry & Technology 28(3): 11–17.

16. Chen YZ, Peng SF, Wang XN (2007) Study of high yield cultivation technologies of oil-Tea *Camellia(Camellia oleifera)*——Formulate Fertilization. Forest research 20(5): 650–655.

17. Carvalho FP (2006) Agriculture, pesticides, food security and food safety. Environmental Science & Policy 9(7-8): 685–692.

18. Watts DB, Torbert HA, Prior SA, Huluka G (2010) Long-term tillage and poultry litter impacts soil carbon and nitrogen mineralization and fertility. Soil Science Society of America Journal 74(4): 1239–1247.

19. Matson PA, Parton WJ, Power AG, Swift MJ (1997) Agricultural intensification and ecosystem properties. Science 277(5325): 504–509.

20. Edmeades DC (2003) The long-term effects of manures and fertilisers on soil productivity and quality: a review. Nutrient Cycling in Agroecosystems 66(2): 165–180.

21. Hernández T, Chocano C, Moreno J, García C (2016) Use of compost as an alternative to conventional inorganic fertilizers in intensive lettuce (Lactuca sativa L.) crops—Effects on soil and plant. Soil and Tillage Research 160: 14–22.

22. Al Seadi T, Drosg B, Fuchs W, Rutz D, Janssen R (2013) 12 - Biogas digestate quality and utilization. The Biogas Handbook: Woodhead Publishing. pp. 267–301.

23. Bond T, Templeton MR (2011) History and future of domestic biogas plants in the developing world. Energy for Sustainable Development 15(4): 347–354.

24. Chen Y (2007)Physiochemical properties and bioactivities of tea seed *(Camellia oleifera)* oil. Dissertations & Theses

25. Liao T, Yuan DY, Zou F, Gao C, Yang Y, Zhang L, Tan X F (2014) Self-Sterility in *Camellia oleiferamay* Be Due to the Prezygotic Late-Acting Self-Incompatibility. Plos One 9(6): e99639.

26. Su MH, Shih MC, Lin KH (2014) Chemical composition of seed oils in native Taiwanese *Camellia* species. Food Chemistry 156: 369–373.

27. Tan C, Ghazali HM, Kuntom A, Tan C, Ariffin AA (2009) Extraction and physicochemical properties of low free fatty acid crude palm oil. Food Chemistry 113(2): 645–650.

28. Vela P, Salinero C, Sainz MJ (2013) Phenological growth stages of *Camellia japonica*. Annals of Applied Biology 162(2): 182–190.

29. He G, Zhang J, Hu X, Wu J (2011) Effect of aluminum toxicity and phosphorus deficiency on the growth and photosynthesis of oil tea (*Camellia oleifera* Abel.) seedlings in acidic red soils. ActaPhysiologiaePlantarum 33(4): 1285–1292.

30. Fan T, Stewart BA, Yong W, Junjie L, Guangye Z (2005) Long-term fertilization effects on grain yield, water-use efficiency and soil fertility in the dryland of Loess Plateau in China. Agriculture, Ecosystems & Environment 106(4): 313–329.

31. Rehl T, Lansche J, Müller J (2012) Life cycle assessment of energy generation from biogas—Attributional vs. consequential approach. Renewable and Sustainable Energy Reviews 16(6): 3766–3775.

32. Hu Y, Cheng H, Tao S (2017) Environmental and human health challenges of industrial livestock and poultry farming in China and their mitigation. Environment International 107: 111–130.

33. Maqbool S, Ul Hassan A, JavedAkhtar M, Tahir M (2014) Integrated use of biogas slurry and chemical fertilizer to improve growth and yield of okra.Science Letters 2(1):56–59.

34. Liu E, Yan C, Mei X, He W, Bing SH, Ding L, Liu Q, Liu S, Fan T (2010) Long-term effect of chemical fertilizer, straw, and manure on soil chemical and biological properties in northwest China. Geoderma 158(3): 173–180.

35. Zheng X, Fan J, Xu L, Zhou J (2017) Effects of Combined Application of Biogas Slurry and Chemical Fertilizer on Soil Aggregation and C/N Distribution in an Ultisol. Plos One 12(1): e170491.

36. Friedel JK (2000) The effect of farming system on labile fractions of organic matter in Calcari-Epileptic Regosols. Journal of Plant Nutrition and Soil Science 163(1): 41–45.

37. Yuan J, Tan X F, Yuan D Y, Zhang X J, Ye SC, Zhou JQ (2013) Effect of phosphates on the Growth, Photosynthesis, and P Content of Oil Tea in Acidic Red Soils. Journal of Sustainable Forestry 32(6): 594–604.

38. Kashem MA, Akinremi OO, Racz GJ (2004) Extractable phosphorus in alkaline soils amended with high rates of organic and inorganic phosphorus. Canadian Journal of Soil Science 84(4): 459–467.

